# Laser scanning identifies large trees as a major source of uncertainty in mangrove carbon accounting

**DOI:** 10.64898/2026.06.12.731900

**Authors:** Toby D. Jackson, Jasper Feyen, Laura Lozano-Arias, JP Caicedo-Garcia, Paula Cristina Sierra-Correa, Cristian Camilo Montes-Chaura, Alicia A. Sanjur, Jorge Hoyos-Santillan, Esperanza González-Mahecha, Diana Castillo, Yuheimy Castillo, Verginia Wortel, Marco P. Ouboter, Noraisah S. Tjong-A-Hung, Danitcijo L. Amiemba, Chantal Rambharos, Consuela Paloeng, Valentien Moe Soe Let, Rashnie Hardin, Faybion Porter, Okieve Kerr, Diego I. Rodríguez-Hernández, Mathilda A. Digby, Tommaso Jucker, Fabian Jörg Fischer, Kim Calders, Charles Price, Farrah Mathura, Hamish Asmath

**Affiliations:** School of Biological Sciences, University of Bristol, Bristol, UK; Q-ForestLab, Department of Environment, Faculty of Bioscience Engineering, Ghent University, Belgium; University of the West Indies, St. Augustine Campus, Trinidad and Tobago; Institute of Marine and Coastal Research of Colombia (INVEMAR); Smithsonian Tropical Research Institute, Panama City, Panama; Landscape and Carbon Dynamics CoLab, School of Forest Sciences, University of Eastern Finland, Joensuu, Finland; Inter-American Development Bank (IDB), Edificio Tower Financial Center, Panama City, Panama; Carbon Laboratory, Center for Hydraulic and Hydrotechnical Research (CIHH), Universidad Tecnológica de Panamá, Panama; Department of Forest Management, Centre for Agricultural Research in Suriname, CELOS, Paramaribo, Suriname; Foundation for Forest Management and Production Control, SBB, Suriname; Forestry Department (Jamaica); Department of Life Science Systems, Technische Universität München, Hans-Carl-von-Carlowitz-Platz 2, 85354 Freising, Germany; Ecology and Evolutionary Biology, University of Tennessee, Knoxville, Tennessee, USA

## Abstract

**Background:** Mangrove forests are crucial ecosystems which support biodiversity, protect coastlines and store vast amounts of carbon. Mangrove conservation and protection rely on accurate carbon accounting to unlock investment. However, the allometric equations underpinning these carbon estimates remain poorly constrained, particularly for the large trees.

**Methods:** We used terrestrial laser scanning (TLS) to estimate the biomass of 187 mangrove stems across Suriname, Panama, Colombia and Jamaica, including 84 stems >20 cm DBH. TLS-derived biomass estimates were used to evaluate local, regional and pantropical allometric equations.

**Results:** Most diameter-based allometric equations underestimated biomass by 8–65%. Equations additionally incorporating tree height performed better, but still underestimated biomass by 12–16% on average. Applying alternative allometries to a representative mangrove inventory from Panama produced biomass estimates ranging from 80 to 200 Mg ha⁻¹, demonstrating that allometric uncertainty alone can generate more than a two-fold difference in estimated carbon stocks.

**Conclusions:** Current allometric equations systematically underestimate the biomass of large mangrove trees and are therefore likely to underestimate mangrove carbon stocks. TLS provides a practical, non-destructive approach for expanding biomass datasets and improving allometric equations. Reducing allometric uncertainty should be a priority for strengthening blue carbon accounting and mangrove conservation.

## Introduction

Mangrove forests are a crucial component of coastal ecosystems and they protect coastlines from extreme weather events (Spalding, Kainuma and Collins, 2010; Zhang *et al*., 2025, 2026). Their combination of high net primary productivity and the exceptional preservation of organic matter in waterlogged, low-oxygen sediments makes them among the most carbon-dense ecosystems on Earth (Donato *et al*., 2011; Zhu and Yan, 2022). However, global estimates of mangrove carbon stocks remain highly uncertain due to limited field data and poor harmonisation of measurement and modelling approaches across scales. A key source of uncertainty arises from the allometric equations used to convert trunk diameter and / or tree height measurements into estimates of aboveground biomass and carbon content (Chave *et al*., 2014). Reducing this uncertainty is therefore essential to ensure that the carbon value of mangrove ecosystems is accurately quantified and effectively protected.

Allometric equations are widely used across tropical forests (Chave *et al*., 2014), but there is a concern that these general allometries do not capture the distinctive architecture and wood properties of mangroves (Kauffman and Donato, 2012). To develop mangrove-specific allometric equations, multiple studies have measured the biomass of mangrove trees in the field by cutting them down and weighing them. However, this is physically challenging and often illegal in mangrove forests. Therefore, mangrove biomass studies usually have a small sample size and focus on developing local, species-specific allometric equations (Fromard *et al*., 1998; Smith and Whelan, 2006; Yepes *et al*., 2016). While these allometric equations may be locally accurate, it is unclear whether they can generalize to the national or regional scale. To address this limitation Price *et al*.(2024, 2026) recently synthesized all available mangrove-specific field measurements of biomass from the Americas and proposed new allometric equations based on either tree height or trunk diameter alone. However, despite the larger sample size (n = 742), this dataset contains very few large trees (12 trees over 30 cm dbh) which results in greater uncertainties for the trees which store the majority of forest carbon (Picard *et al*., 2025). Furthermore, we expect that large tree allometries may differ systematically from those of small trees because they are more exposed to the elements once they reach the canopy.

One solution to estimate the biomass of trees without cutting them down is to use terrestrial laser scanning (TLS). TLS produces detailed 3D point cloud representations of the trees, from which the total volume of wood in the trunk and branches can be estimated (Disney *et al*., 2018). The biomass of the tree is then given by multiplying the wood volume by the wood density. For wood density, most studies use species-specific data from the literature or measurements from local conspecifics (Phillips *et al*., 2019), although species wood density can vary with environmental factors (Fischer *et al*., 2026). This combination of TLS and wood density measurements has been shown to produce accurate biomass estimates for diverse forest types ranging from Eucalypt forests in Australia (Calders *et al*., 2015), tropical forests (Gonzalez de Tanago Menaca *et al*., 2017) and conifer plantations (Yrttimaa *et al*., 2022). The TLS method can provide a larger and more representative sample of trees available for allometric equations. For example, recent work used TLS to show that the biomass of temperate woodlands in the UK had been underestimated by 75 % by previous allometric equations (Calders *et al*., 2022) or by up to 40% in tropical forests (Terryn *et al*., 2024).

This study was part of an Inter-American Development Bank project aiming to harmonize mangrove carbon monitoring across Suriname, Panama, Colombia, and Jamaica. Mangrove measurement protocols in each country currently recommend site-specific allometric equations based on limited sample sizes. In order to test their accuracy, we estimated the aboveground biomass and carbon storage of 187 mangrove stems using TLS, focusing on the large stems which dominate forest carbon storage (84 stems over 20 cm DBH and 33 stems over 30 cm DBH). We exclusively scanned *Avicennia germinans* and *Rhizophora* spp., which are the dominant mangrove species in the region (Price *et al*., 2024). We use these TLS biomass estimates to test whether current allometric equations, either local and regional, accurately predict the biomass of large mangrove trees.

## Methods

### Description of the study sites

This study sampled 187 mangrove trees in Suriname, Colombia, Panama and Jamaica.

In Panama, mangroves are distributed across the Pacific (97%) and Atlantic (3%) coasts, covering 1,870 km². The dominant species are *Rhizophora mangle* L. and *Rhizophora racemosa* G. Mey., which occur in both monospecific and mixed forest stands. Other species, including *Conocarpus erectus* L., *Avicennia germinans* (L.) L., *Avicennia bicolor* Standl., *Laguncularia racemosa* (L.) C.F. Gaertn., and *Pelliciera rhizophorae* Planch. & Triana, are also found as monospecific or mixed stands, with hydrogeomorphic settings strongly influencing forest composition and the spatial distribution of species. Mangroves in Panama experience contrasting climatic regimes. On the Pacific coast, the climate is characterized by two wet seasons (May–July and August–November) and two dry seasons (December to April and July-August), with annual precipitation ranging from 1,500 to 3,500 mm y^-1^. On the Atlantic coast, rainfall occurs year-round, exceeding 4,000 mm y^-1^. Panama has two distinct, contrasting tidal regimes on each of its coasts. On the Pacific, Panama experiences a high-range, semi-diurnal tide; while on the Atlantic, a mixed tidal regime oscillating between semidiurnal and diurnal has a very low range.

In Suriname, the national mangrove area was estimated at 876 km², consisting of approximately 70% coastal *Avicennia*-dominated and 30% *Rhizophora* dominated mangroves. The country has a tropical climate with mean annual temperatures around 32 °C and an average annual precipitation of 2200 mm, characterized by two wet seasons (April–August and November–February) and two dry seasons (August–November and February–April). Suriname’s coastline is further shaped by a semi-diurnal microtidal regime, with two high and two low tides per day and a mean tidal range of 1.8 m, increasing to 2.5 m during spring tides.

In Colombia, mangroves cover approximately 2750 km², of the coastline, with 71 % on the Pacific and 29 % on the Caribbean coasts. Data collection for this study took place in the “Mangrove and Lagoon Ecosystem Ciénaga de la Caimanera”, a coastal lagoon on the Caribbean coast. This study site is part of the reference area for the carbon estimation certification project known as Vida Manglar in the Gulf of Morrosquillo. Due to its spatial extent, structural characteristics, and heterogeneous conservation status, this area represents a relevant setting for evaluating carbon accumulation potential in the the three tree species present, *Rhizophora mangle*, *Avicennia germinans* and some individuals of *Laguncularia racemosa*, and generating technical information to strengthen the blue carbon baseline for mangroves in the Colombian Caribbean. The “Cienaga de La Caimanera” is characterized by a bimodal precipitation regime, with a mean annual rainfall of 1263 ± 415.8 mm, reaching minimum values in January (6.64 mm) and maxima in July (171.1 mm). Mean air temperature ranged between 27.0 °C in October and 28.9 °C in March, while relative humidity averaged 81.1 ± 0.4% and mean wind speed was 1.97 ± 0.06 m*s-1. Historical tidal amplitude reached approximately 1.0 m (Caicedo-Garcia *et al*., 2025).

In Jamaica, mangrove cover was approximately 138 km² prior to the passage of hurricane Melissa in October, 2025, of which over 20 km² were impacted by the hurricane across several parishes. Over 80% of Jamaica’s mangrove habitats are located on the southern coastline of the island. Data collection was conducted across mangrove ecosystems located along the southern coastline of Clarendon within the broader Portland Bight region and associated coastal wetland systems. This region represents one of Jamaica’s most ecologically significant mangrove landscapes, containing extensive mangrove forests distributed between the Milk River and Salt River coastal systems. Over recent decades, these coastal systems have experienced substantial degradation due to storm impacts, altered hydrological regimes, charcoal production, and other anthropogenic pressures. Consequently, restoration initiatives have identified the southern Clarendon coastline as a priority area for ecosystem rehabilitation and blue carbon conservation. The study area consisted of a mixed mangrove system dominated by *Rhizophora mangle*, *Avicennia germinans*, and *Laguncularia racemosa*, occurring within interconnected wetlands, tidal channels, and shallow coastal environments. Jamaica experiences a tropical maritime climate characterized by warm temperatures throughout the year, with mean annual temperatures generally ranging from 26–28 °C and annual precipitation across the southern coastal plain ranging between approximately 800 and 1,500 mm. Rainfall patterns are influenced by bimodal wet seasons typically occurring between May–June and September–November, separated by relatively drier periods. Coastal hydrology along southern Jamaica is influenced by a microtidal regime with relatively small tidal amplitudes, seasonal freshwater inflows, and periodic exposure to tropical storms and hurricanes, all of which contribute to the structural and ecological dynamics of mangrove ecosystems within the Portland Bight area.

### Terrestrial laser scanning data collection

In Suriname, a RIEGL VZ-400i terrestrial laser scanner was used with an angular resolution of 0.04 degrees and a scanning frequency of 600 kHz. We surveyed plots sized 20 × 50 m using a grid-based scanning pattern, with approximately 10 m spacing between scan positions, resulting in 18 scan locations per plot. At each location, an additional scan was collected with the instrument tilted 90° from the vertical to minimize occlusion due to the restricted zenith range of the instrument. This approach produced 36 scans per plot, ensuring dense coverage of tree stems and canopy structure. All scans were co-registered in RIEGL’s RiSCAN Pro software to generate plot-level point clouds. Registration was further refined using the Multi Station Adjustment Algorithm, in which the orientation and position of each scan position were iteratively adjusted to minimise the global fitting error of tie-planes - planar surfaces shared across overlapping scans. From the resulting plot-level point clouds, individual trees were manually identified and segmented for analysis.

In Panama, Colombia and Jamaica individual trees were scanned in a targeted approach using a Leica BLK 360 G2 Laser Scanner (G2 Survey, UK). Tree selection targeted large individuals of common species, because these are poorly sampled in existing allometric analyses. Given the lower quality of the Leica scanner, compared to the RIEGL, we adjusted the sampling design to maximize the quality of the resulting point cloud and increase the comparability of the two data sets. The target tree was scanned from multiple positions (25 to 60) arranged in concentric circles around it. Targets were fixed in place throughout the scanning process and used to combine the multiple scans into a single point cloud.

The point clouds were aligned and the initial cleaning of unwanted material was done with Cyclone Register 360 plus software from Leica Geosystems. All tree point clouds were checked for data quality issues such as blurring of the point cloud due to wind, or undersampling of the upper branches. Trees with clear data quality issues were removed at this stage.

### Terrestrial laser scanning data cleaning

The quality-controlled tree point clouds were then manually cleaned in CloudCompare. This involved manually removing all the leaves, noise and points belonging to neighbouring trees from the point cloud. This labour-intensive data cleaning is the most reliable way to produce high-quality tree-level point clouds. It is also a subjective process, where the individual cleaning the data must regularly decide whether a particular set of points represents a branch or noise. As the point density reduces near the top of the canopy it becomes increasingly difficult to distinguish branches from noise, so most of our tree-level point clouds are missing these small branches. All tree-level point clouds, before and after cleaning, are made openly available online so readers can assess data quality. These manually cleaned point clouds could also serve as a valuable benchmark for assessing the accuracy of automatic data cleaning algorithms.

Where a single tree split into multiple stems below 1.3 m, we treated each stem separately in line with protocols (S1). We split the TLS point cloud data into separate files for each stem, modelled their biomass separately and included them separately in the analyses below. In some cases, multi-stemmed trees grow from a single set of stilt roots. Field plots measure each stem individually. We therefore treated them individually for the purpose of developing allometric equations.

We measured the DBH and height of each stem on the point cloud using the measuring tool in CloudCompare and following the field protocols for mangrove trees (see S1). The height was simply measured as the vertical distance from the lowest point to the highest point that clearly belonged to this stem. Height measurements are notoriously difficult in a dense forest so our point cloud based measurements are likely to be more accurate than field estimates. By default, we measured the DBH at 1.3 up the stem. However, most of the *Rhizophora* trees in our sample had stilt roots and in most cases we measured DBH just above the highest root. The DBH was measured by cutting a cross section of the trunk in CloudCompare (Fig 1) and measuring its diameter in two directions and then taking the mean (see table 1).

**Figure 1.**
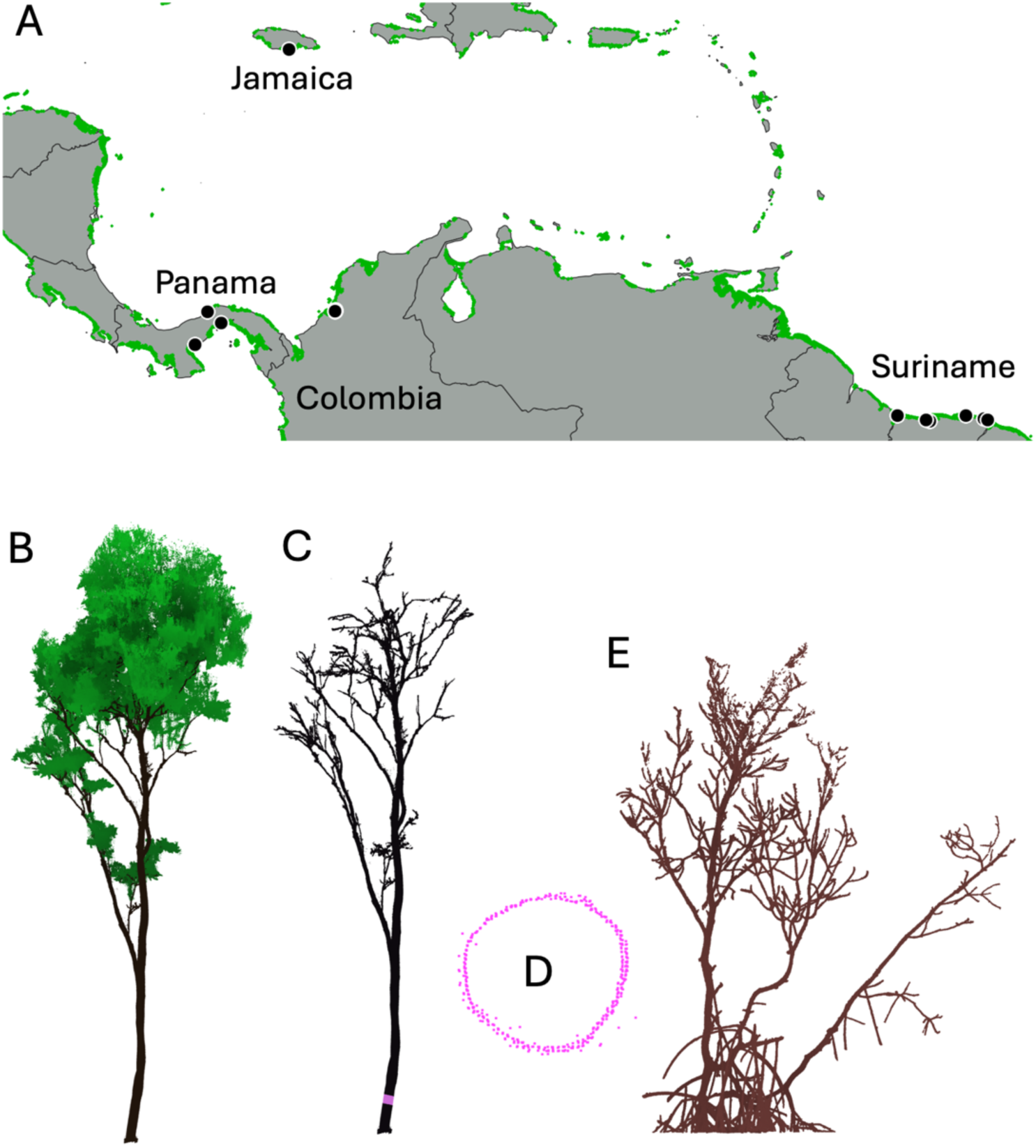
A: Map of sampling locations (points) across four countries with the mangrove extent shown in green. B: Example TLS point cloud for a 30 m tall Avicennia racemosa in Suriname with manually separated leaves coloured green. C: The same tree without leaf points, showing the level of branching detail available for volume estimation. D: Cross section of the trunk point cloud used to measure DBH. E: Example leaf-off point cloud for a 12 m tall Rhizophora mangle in Jamaica with three stems and stilt roots.

**Table 1.** Sample sizes and DBH ranges of the trees scanned in each country.

|  |  | Suriname | Panama | Colombia | Jamaica |
| --- | --- | --- | --- | --- | --- |
| *Avicennia germinans* | *N stems* | 49 | 5 | 12 | - |
|  | *Min DBH (cm)* | 5.0 | 20.1 | 14.5 | - |
|  | *Mean DBH (cm)* | 23.3 | 23.9 | 26.5 | - |
|  | *Max DBH (cm)* | 42.6 | 28.9 | 51.7 | - |
| *Rhizophora spp.* | *N stems* | 27 | 27 | 24 | 43 |
|  | *Min DBH (cm)* | 5.3 | 3.0 | 6.8 | 4.9 |
|  | *Mean DBH (cm)* | 20.4 | 21.4 | 23.1 | 13.4 |
|  | *Max DBH (cm)* | 41.2 | 49.6 | 44.3 | 21.4 |

### Estimating woody volume with cylinder models

The total woody volume of each stem was estimated by fitting cylinder models to the point clouds using TreeQSM (Åkerblom, 2017). Prior to cylinder model fitting, the above-ground stilt roots were separated from the original point cloud and inverted to look like upward-branching trees (Asmath *et al*., 2025). The accuracy of the cylinder models depends on the input parameters. To ensure an accurate model, we followed the guidelines and example code provided in the TreeQSM documentation. We started by generating 40 cylinder models covering a wide range of input parameters. From these, we selected the optimal input parameters as the ones which produced the cylinder model with the lowest mean distance between the cylinders and the original point cloud. To account for the inherent randomness of the cylinder fitting process, we then generated 20 cylinder models using the same input parameters. These 20 cylinder models were then used to calculate a mean and standard deviation for the woody volume of each stem. This whole process was conducted separately for the stems and stilt roots, and the woody volumes combined. A common problem is that the small branches are poorly resolved in the point cloud and are therefore difficult to fit cylinder models for (Demol *et al*., 2022; Morales and MacFarlane, 2025). We tested the sensitivity of our cylinder models to the small branches by comparing the total woody volume with and without all branches below 5 cm diameter (S2). We also repeated the cylinder fitting process using an independent method (RayCloudTools) and compared the results (S3).

Once the woody volume has been estimated from TLS, the biomass (in kg) was calculated by multiplying this volume by the wood density derived from the Global Wood Density Database (Fischer et al 2026). For the purpose of our analysis, we combined *R. mangle* and *R. racemosa* because they are difficult to distinguish in the field in the absence of inflorescences and they usually have similar wood densities.

### Measuring wood carbon content

Where possible, wood cores were collected from the scanned trees for analysis of carbon content. Wherever possible, wood cores were collected from target trees using an increment borer 3-Thread, 0.200 (5.15mm) (Jim-Gem, Forestry Suppliers, USA). However, in some cases we extracted a rectangular section of wood approximately 3 x 3 x 1 cm in size. Wood samples were dried at 80 **°** C in a Fisher Scientific Isotemp oven and finely ground in a ball mill. The carbon content was then measured using a Thermo Flash EA 1112 elemental analyzer. We note that wood cores were not collected from the same trees that were scanned in Suriname and Panama because the scanning took place before this project started.

### Assessing allometric equations

We used the TLS-derived biomass estimates to test the validity of three types of published allometric equations (Table 2). First, we tested local allometric equations predicting biomass from species and DBH which are currently recommended to estimate mangrove aboveground biomass in each project country. For Suriname, we tested an equation based on mangroves from French Guiana (Fromard *et al*., 1998). For Jamaica, we tested an equation based on mangroves from the Florida Everglades (Smith and Whelan, 2006). For Colombia and Panama, we tested an equation based on mangroves from Colombia (Yepes *et al*., 2016). Secondly, we tested the regional equation for predicting mangrove biomass from DBH, but not species, based on a collated data set of 630 destructively harvested mangroves across the Atlantic East Pacific (AEP) biogeographic region (Price *et al*., 2026). Finally, we tested three allometric equations which include tree height and wood density as well as DBH to predict biomass. Two of these are mangrove specific (Chave *et al*., 2005; Price *et al*., 2026) while the third is recommended for all tropical trees Chave et al. (2014). We applied a Baskerville correction when transforming from models fit in log-space to actual biomass predictions. However, this was not possible in Suriname and Jamaica because the residual errors were not reported. These errors are usually 1-2 %, so unlikely to influence our conclusions.

**Table 2.** Allometric equations tested in this study.

| Reference | Species (N) | Equation |
| --- | --- | --- |
| Fromard et al (1998) | Avicennia (25) | $Biomass = 0.14 \times DBH^{2.4}$ |
| | Rhizophora (9) | $Biomass = 0.1282 \times DBH^{2.6}$ |
| Smith & Whelan (2006) | Avicennia (8) | $\log_{10}(Biomass) = 1.934 \times \log_{10}(DBH) - 0.395$ |
| | Rhizophora (14) | $\log_{10}(Biomass) = 1.731 \times \log_{10}(DBH) - 0.112$ |
| Yepes et al (2016) | Avicennia (30) | $\ln(Biomass) = 2.45 \times \ln(DBH) - 1.96$ |
| | Rhizophora (30) | $\ln(Biomass) = 2.59 \times \ln(DBH) - 1.91$ |
| Price et al (2026) | Mangroves in the AEP (630) | $\ln(Biomass) = 1.982 \times \ln(DBH) - 0.956$ |
| | | $\ln(Biomass) = 0.801 \times \ln(Height \times \rho \times DBH^2) - 1.482$ |
| Chave et al (2005) | Mangroves (84) | $Biomass = 0.0509(Height \times \rho \times DBH^2)$ |
| Chave et al (2014) | 4004 tropical trees | $Biomass = 0.0673(Height \times \rho \times DBH^2)^{0.976}$ |

The crucial question is whether the plot-level biomass is systematically under or over predicted, so we decided to use the average bias expressed as a percentage of the biomass in the sample. The use of the bias is particularly valuable as our dataset is approximately representative of the biomass distribution found in the average mangrove forest plot in these regions (i.e. more large trees). These bias estimates therefore approximate the errors we would expect when using these equations for estimating plot biomass, which is their intended purpose. We also calculated the coefficient of determination (R^2^) and found very high correlations (R2 > 0.9) even where allometric equations visually performed quite poorly. We also tested the root mean square error (RMSE) and found high RMSE for all equations (60-90%). However, high RMSE is not necessarily a problem if the errors are randomly distributed, because they will cancel out at the plot-scale where these equations are used (Picard *et al*., 2025).

### Implications for plot-level biomass stocks

To demonstrate the implications of allometric equations and uncertainty on plot level biomass stocks, we used the openly available plot inventory data for mangroves in Panama (Hoyos-Santillan, Miranda, *et al*., 2025). This data set contains DBH measurements for 453 *Avicennia* and 951 *Rhizophora* trees across a network of plots. We compare the recommended allometric equation (Yepes *et al*., 2016) with the regional equation (Price *et al*., 2026) and the pantropical tree equation (Chave et al 2014). We also compare a third allometric equation based on the TLS biomass data presented in this study. For this equation, we fit a linear model to predict log-transformed biomass from log-transformed DBH. For each 10 cm DBH size class, we estimated the mean tree biomass using each allometric equation. We then multiplied these tree-level biomass estimates by the number of trees in that size class to give the total biomass for each size class.

## Results

### Carbon content of mangrove trees

The mean carbon content was 46.7 ± 1 % across all 110 trees sampled in this study. There were statistically significant differences between countries and species (Figure 2). However, the variations in carbon content were far smaller than the variations in biomass described in the following sections. We therefore focus on biomass variation, on the understanding that approximately 46.7 % of this biomass is carbon.

**Figure 2.**
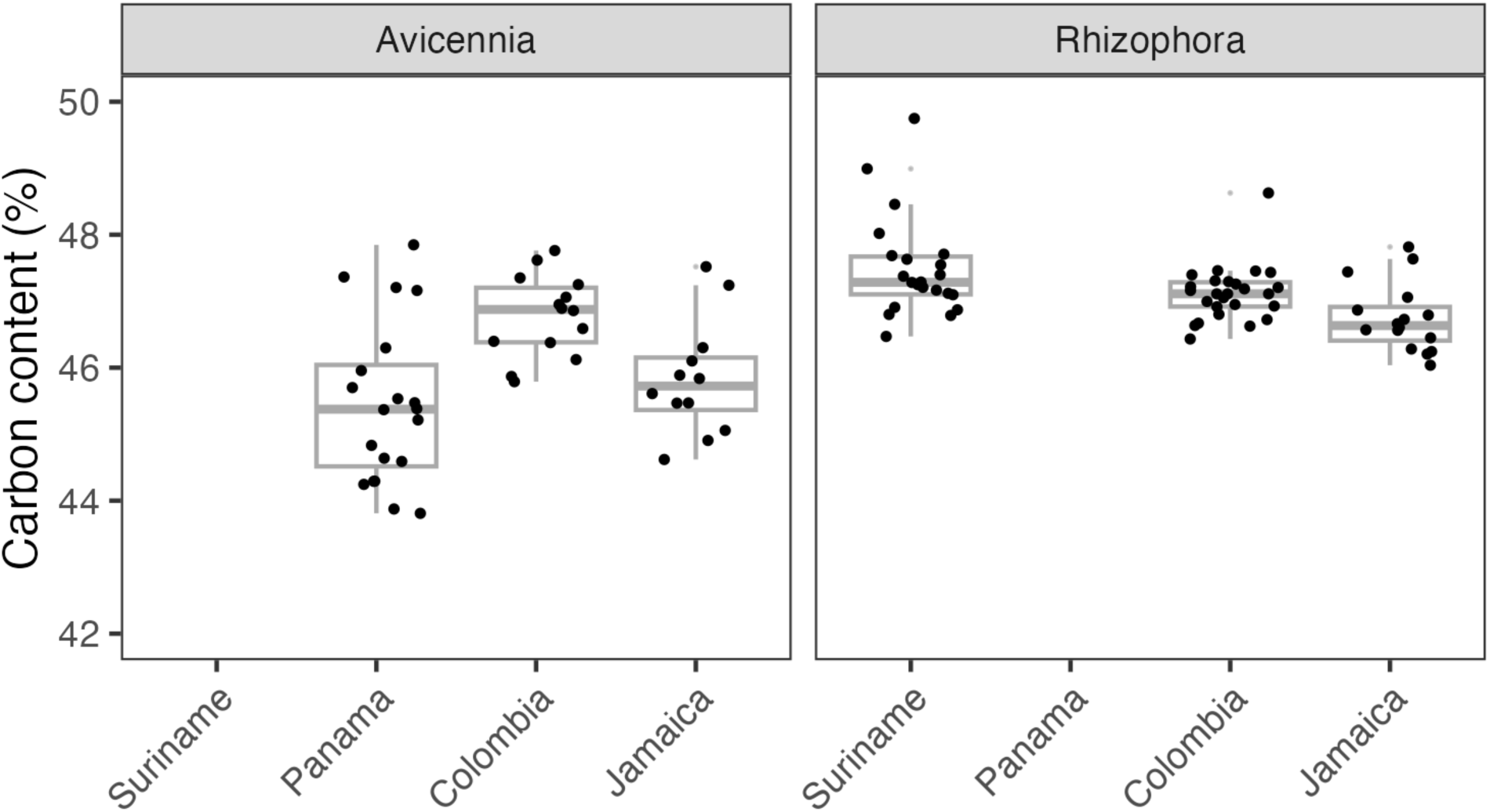
Carbon content (%) of the wood cores sampled in this project.

### Biomass estimates from terrestrial laser scanning

Our TLS-derived biomass estimates ranged from 4 kg to 5276 kg per mangrove stem, with a mean of 513 kg (Figure 3). The standard deviation was 13 % of the biomass and did not vary with tree size. For a given DBH, *Rhizophora* had consistently higher biomass than *Avicennia*, largely due to their stilt roots which contained 29 % of their biomass on average. Mangroves from Suriname, Colombia and Panama spanned almost the entire biomass range, whereas those from Jamaica were considerably smaller (22 - 328 kg). This reflects a biogeographical difference where the mangroves in Jamaica tend to be short and multi-stemmed, perhaps due to the periodic hurricanes in this region.

**Figure 3:**
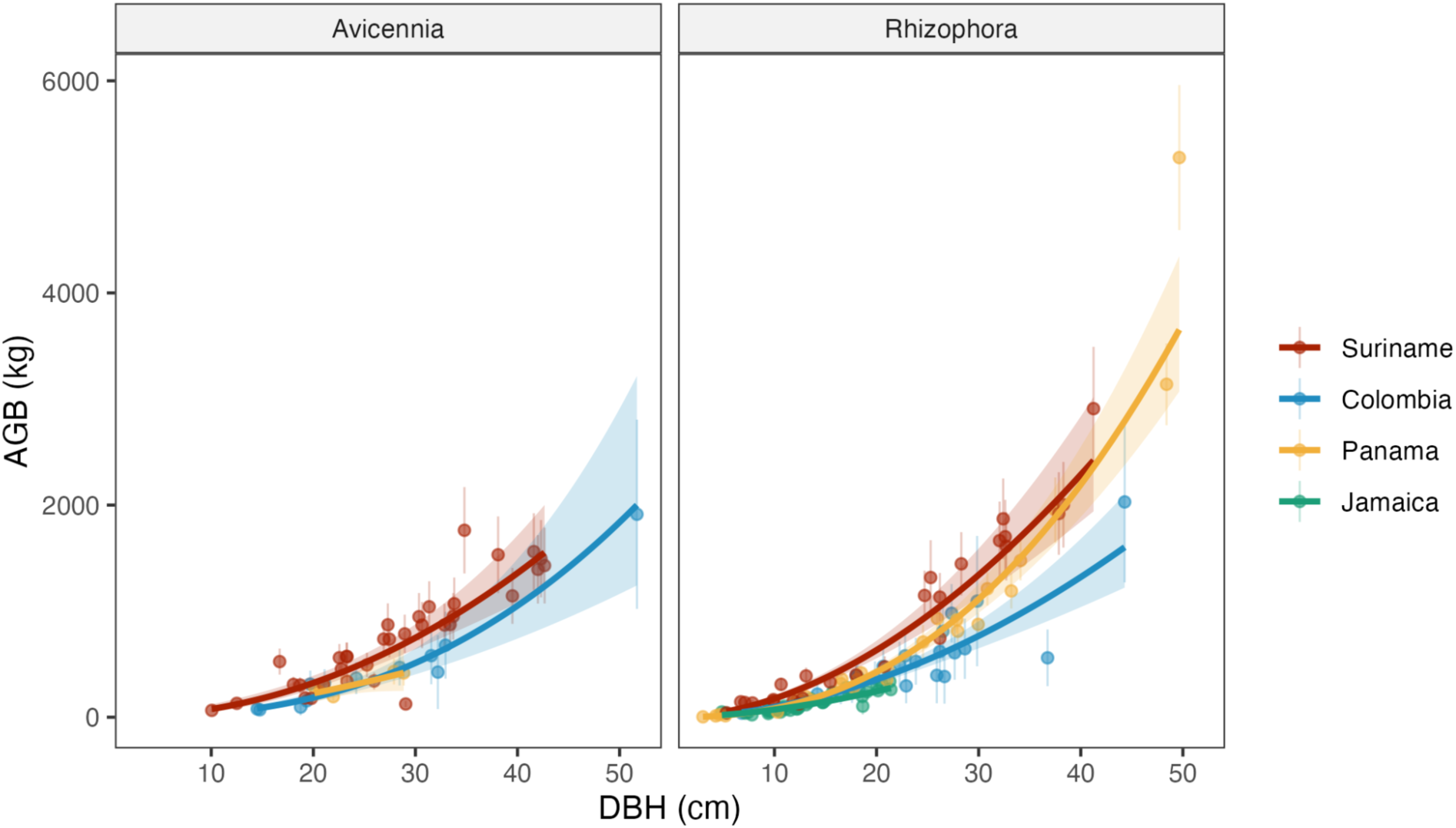
Biomass allometries for trees in this study subdivided by country and genus.

### Comparing local and regional diameter-based allometries

All diameter-based allometric equations underestimated mangrove biomass by 8 to 65 % (Figure 4). The only exception was the Yepes et al. (2016) equation, which overestimated biomass by 6 % in Colombia (Table 3). Locally recommended equations performed significantly better than the regional equation of Price et al (2026) in Suriname, Panama and Colombia (see table 3). This is likely because these local equations were based on trees growing in similar climatic conditions (French Guiana for Fromard et al. (1998) and Colombia for Yepes et al. 2016). However, the regional equation performed better than the locally recommended equation in Jamaica, underestimating biomass by 35 % as opposed to 44 %.

**Figure 4.**
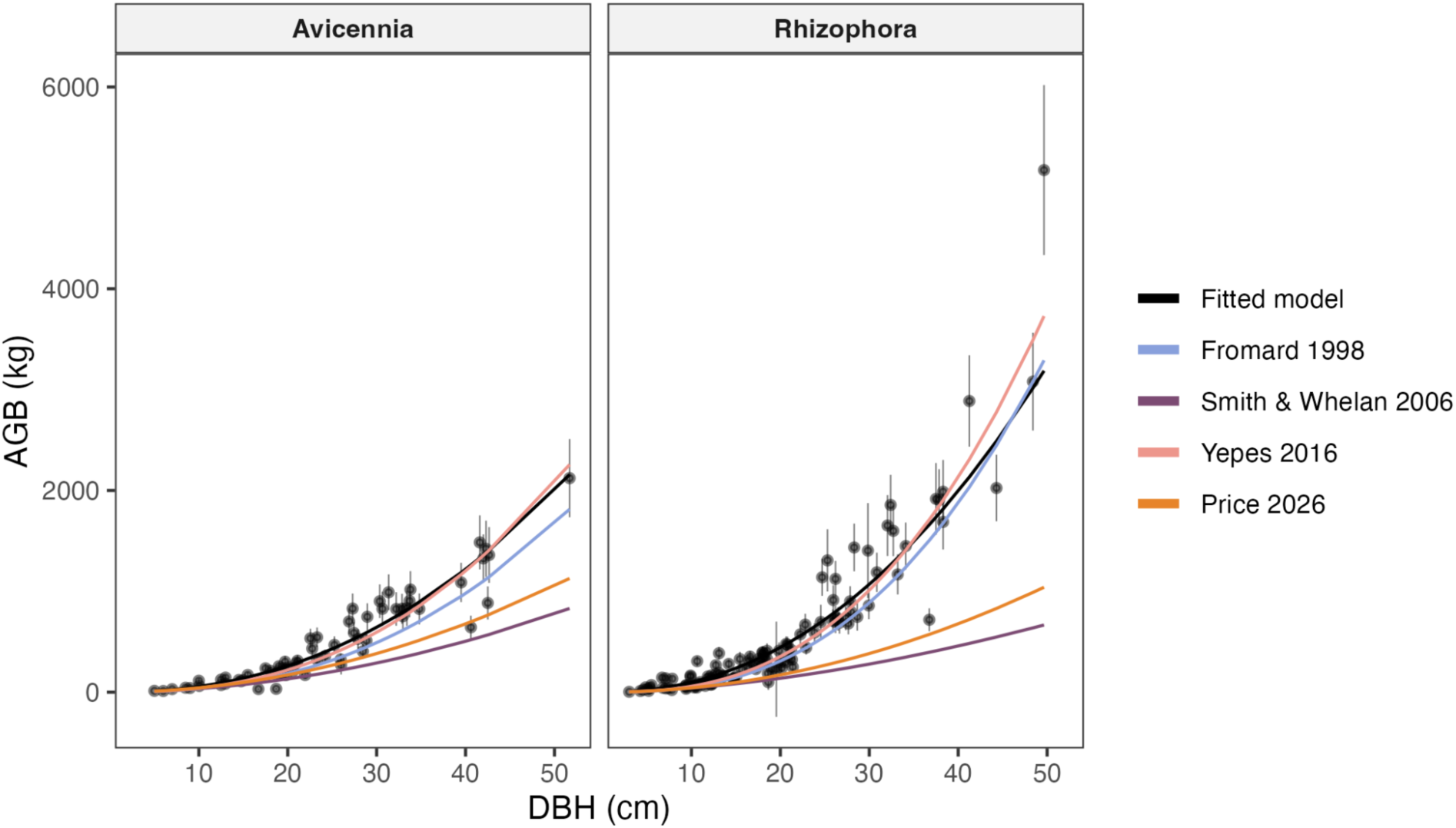
Testing diameter based allometric equations. The points represent the TLS biomass estimates while the coloured lines represent existing allometric equations. An identical figure on logarithmic scales is given in Figure S4 and the residuals are plotted in Figure S5.

**Table 3.**
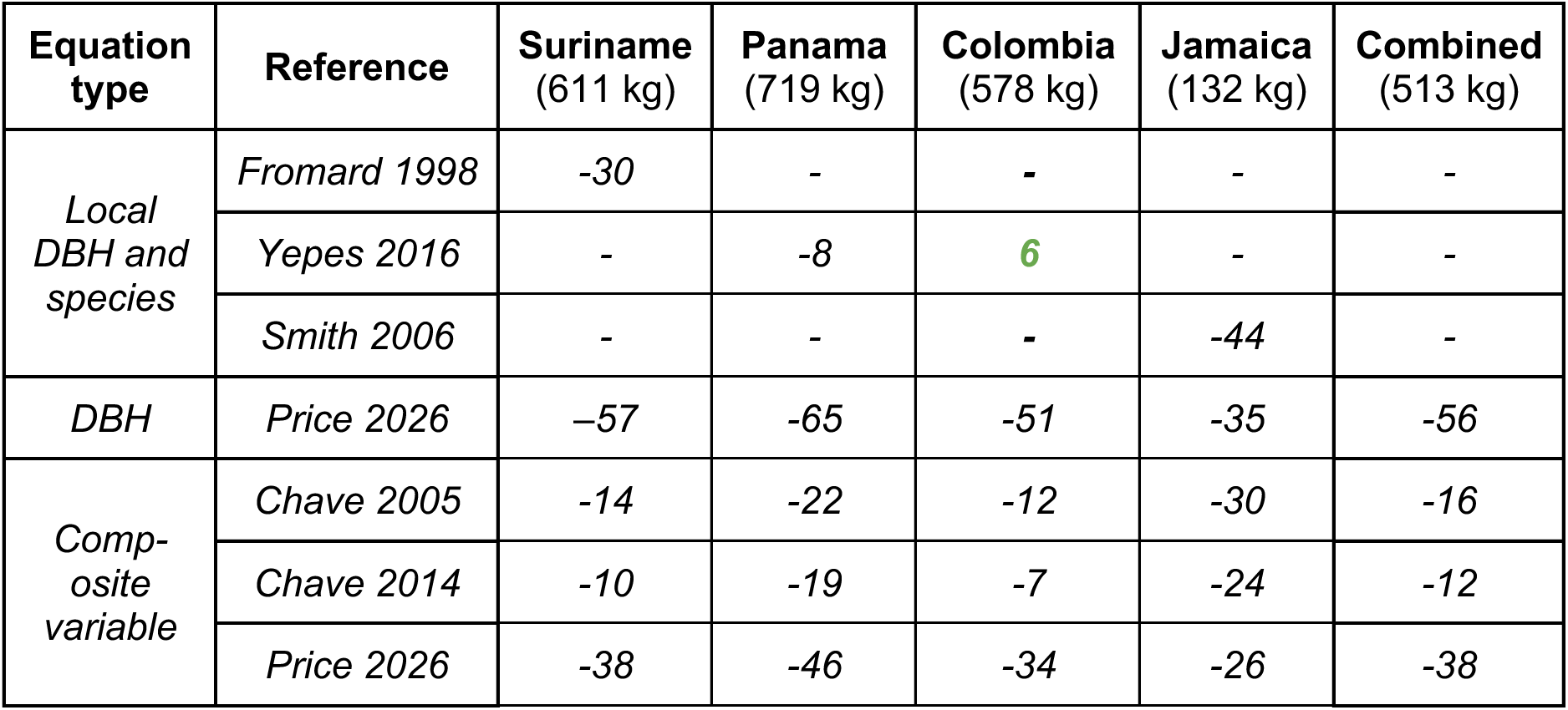
Bias of allometric equations assessed against TLS derived biomass estimates. Bias is expressed as a percentage of the mean tree biomass per country. Note that all values are negative except the single entry highlighted in green.

### Accuracy of allometries including height and diameter

As expected, we found that allometric equations including height measurements as well as DBH generally outperformed those based on DBH only. We found that the mangrove-specific allometry of Chave et al. (2005) was very similar to the pantropical tree allometry of Chave et al. (2014). These allometric equations underestimated the overall mangrove biomass by 16 % and 12 % respectively (Figure 5). The underestimation was modest in Colombia, Suriname and Panama, but considerably larger in Jamaica (30 % and 24 %, respectively). In contrast, the Price et al. (2026) allometry significantly underestimated across all four countries by between 26 and 46 %.

**Figure 5:**
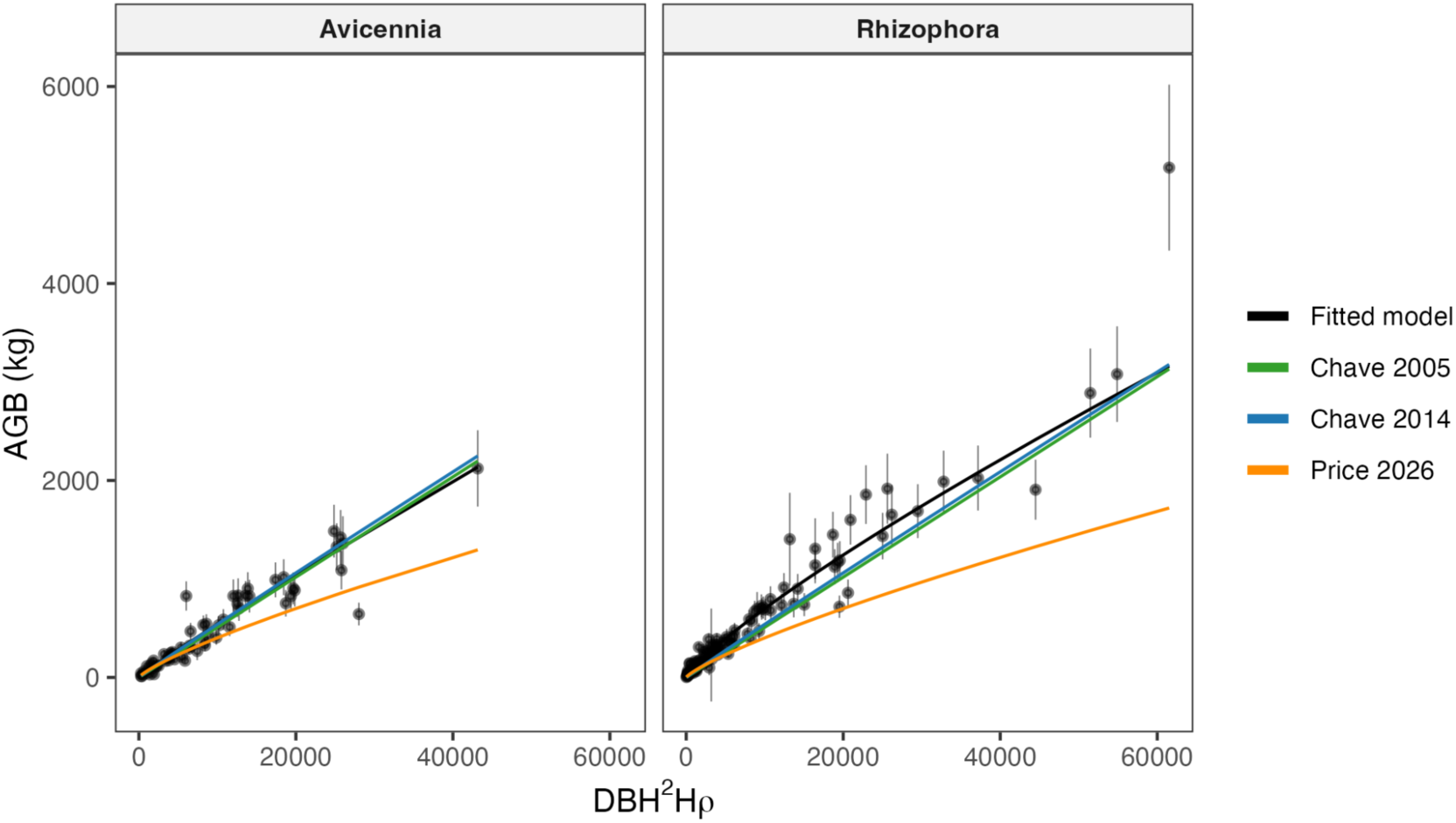
Testing allometric equations based on a composite variable containing tree height, diameter and wood density (fixed for each species). An identical figure on logarithmic scales is given in Figure S4 and the residuals are plotted in Figure S5.

### Implications for plot-level biomass stock estimates

To demonstrate how uncertainty in allometric equations translates into plot-level biomass uncertainty, we applied four allometric equations to an openly available mangrove inventory data set from Panama (Hoyos-Santillan, Miranda, *et al*., 2025). On average, these forest plots contained 343 trees with a DBH less than 20 cm, compared to 138 with a DBH > 30 cm (Figure 6a). However, each large tree contains substantially more biomass than each small tree. Large trees also have a greater biomass uncertainty, as demonstrated by the difference between plausible allometric equations (Figure 6b). For example, for a tree with a 15 cm DBH the biomass predictions range from 92 kg to 173 kg. In contrast, for a tree with 35 cm DBH the biomass predictions can range from 508 kg to 1462 kg. These tree-level biomass estimates and associated uncertainties are then translated into plot-level biomass uncertainty by multiplying by the the abundance of the trees in each size class (Figure 6c). We see the largest biomass contributions, and the largest biomass uncertainty, derives from trees above 30 cm. Overall, the mean biomass estimates for Panama’s mangroves ranges from 80 Mg ha to 200 Mg ha, depending on the allometric equation. We note that in this illustrative example we removed trees below 5 cm and above 50 cm, because very few of these were recorded in the Panama data set.

**Figure 6.**
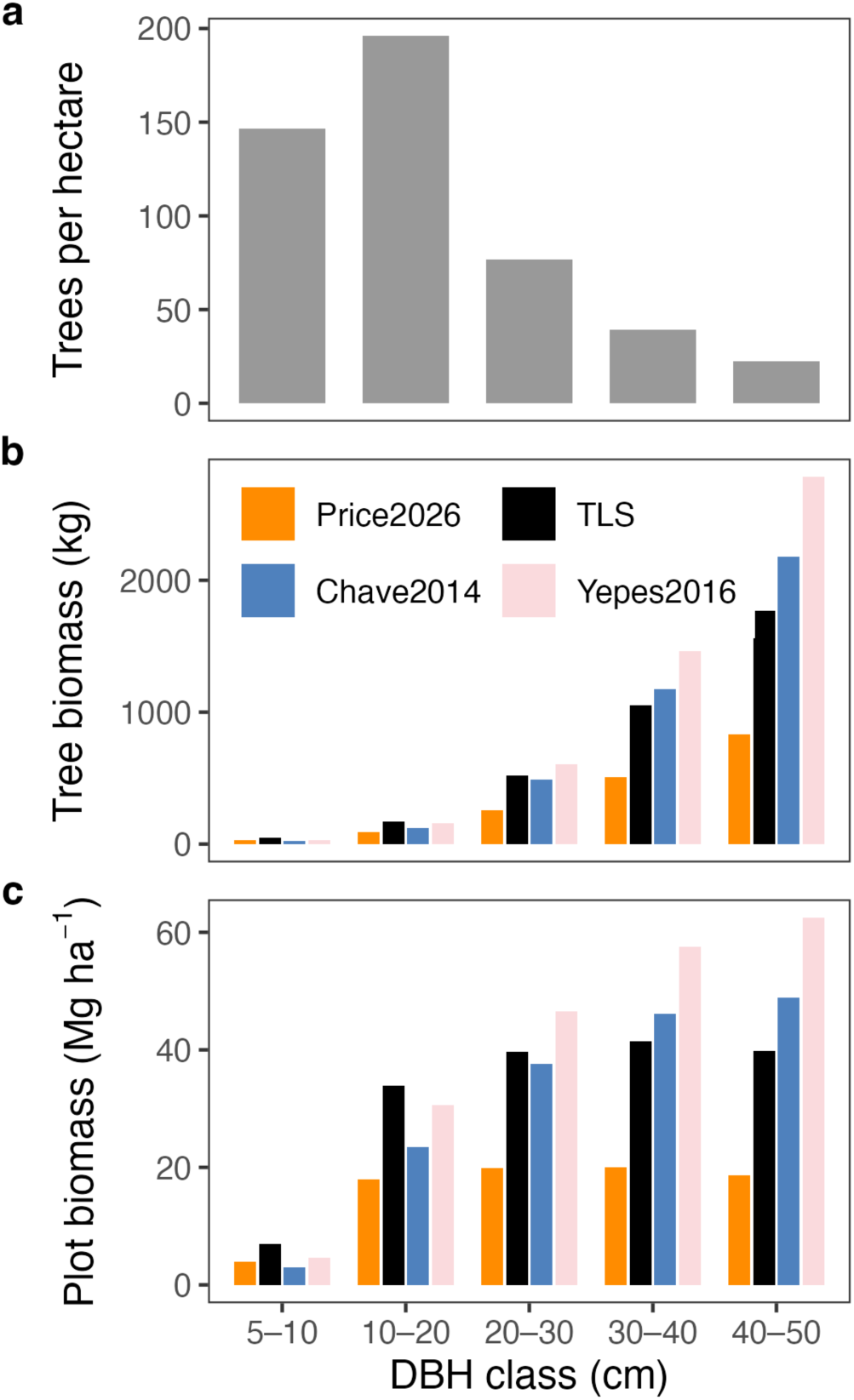
Example of the impact of allometry on biomass estimates across mangrove tree size classes. (a): Number of trees in each DBH size class. (b): Predicted biomass of the mean tree in each size class from four different allometric equations. (c) Contribution of these trees to the plot- level biomass (c = a x b). Note that the first diameter range is smaller than the others because trees < 5 cm DBH are not usually measured.

## Discussion

### Mangrove carbon underestimated by allometric equations

Our results suggest that mangrove forests carbon stocks are higher but also more uncertain than previously understood. This is because current allometric equations underestimate the biomass of mangrove trees, particularly the large trees (>30 cm DBH) which store the most carbon. However, the biomass data for these large trees is so scarce that current allometric equations vary threefold in their plot level biomass estimates. By improving plot-level biomass estimates, regional carbon maps can more accurately capture carbon storage, strengthening the case for protecting and restoring these vital ecosystems.

The scale of this underestimate varies between regions and allometric equations. Pooling all the data in this study (187 stems from four countries: Suriname, Panama, Colombia and Jamaica) we find that the Chave et al. (2014) pantropical allometry underestimates mangrove carbon by 12 %, whereas the Price et al. (2026) regional mangrove allometry underestimates mangrove carbon by 56 %. These large differences between allometric equations underlines the uncertainties in the carbon monitoring pipeline. For example, we reviewed the studies estimating mangrove biomass in the past decade (2015-2025) and found that approximately 30% use Chave et al. (2005 or 2014) while the remaining 70 % generally use more local equations which are likely to be highly variable in accuracy (see S6).

The challenge of building regional or global allometric equations was demonstrated in this study by the Jamaican mangroves, which were smaller and more structurally complex than those in Suriname, Colombia or Panama. Jamaican mangroves were commonly multi-stemmed, with up to seven stems per tree. Furthermore, they generally had large stilt root systems which comprised a significant proportion of their biomass, up to 60 % in some cases. In this study, we decided to treat each stem individually, measuring its DBH, height and biomass separately from its neighbours. The biomass of the stilt roots was then split between the stems based on connectivity and proximity. However, replicating this consistently in field surveys may prove challenging. One of the benefits of TLS is that the 3D model of the trees are permanently available, meaning that future researchers can test different sampling strategies for complex cases like these.

### Robust biomass estimates from terrestrial laser scanning

We ensured that our TLS-derived volume estimates were robust by carefully optimizing the data collection, cleaning and processing. (1) We designed our data collection to maximize data quality, targeting specific trees with as many as 60 scan positions using the Leica scanner. (2) We then manually cleaned each individual tree, removing all the leaves and any branches from neighbouring trees. This manual data cleaning process was extremely labour intensive but necessary to prepare the point clouds for the cylinder fitting step. (3) We tested a range of parameters for the cylinder fitting process, selecting the optimal cylinder model for each tree by minimizing the point-to-cylinder distance. (4) Finally, we tested the sensitivity of our results by running two sensitivity analyses. We demonstrated that our tree volume estimates were not sensitive to the treatment of small branches (S2), which are often poorly modeled due to poor data coverage in the upper canopy (Demol *et al*., 2022; Morales and MacFarlane, 2025). We compared our results to an independent cylinder fitting algorithm and found a strong agreement (R^2^ = 0.89, bias = 0.029 m^3^, see S3). We are therefore confident that our tree volume estimates are robust and we make all of the data openly available for others to confirm.

To calculate tree biomass, we multiplied the tree volume by the wood density. There is therefore the potential for uncertainty or bias in the wood density measurements to impact our biomass estimates. We tested both local and global wood density estimates and found that, while there was a significant difference between them, the impact on our results was relatively small. However, one potential issue with all of the wood density measurements is that they are based on wood cores taken from the lower part of the trunk (i.e. where a person can easily reach). Therefore, if wood density varies systematically with tree height (Olagoke *et al*., 2016) this single-point measurement will be biased.

### Future research priorities

Our results emphasize the importance of including large and medium sized trees in biomass allometries. More biomass measurements from large destructively harvested trees would therefore be highly valuable. We emphasize that these destructive harvest data are extremely difficult to collect. To maximize their value we therefore suggest that, wherever possible, trees should be carefully scanned with a high quality terrestrial laser scanner before being cut down. This will allow researchers to improve the robustness of TLS-based approaches and therefore reduce the need to cut down mangrove trees in the future.

We decided not to propose new allometric equations in this study because, despite the inclusion of large mangrove trees, we had a relatively low sample size for each study site. However, we strongly support efforts to develop new allometric equations by collating existing mangrove biomass data (for example see Price et al. 2024 and 2026) and our data is openly available for this purpose. Increased data sharing and synthesis efforts will help reduce uncertainty in allometric equations and improve our estimates of mangrove carbon storage. We recommend that synthesis efforts include both destructive harvest data and TLS-based biomass estimates and build models which consider local environmental factors such as tidal range, salinity and the disturbance regime. One productive way forward could to use global data sets to model the variation of tree height (Jucker *et al*., 2022) and wood density (Fischer *et al*., 2026) with environmental factors and then predict biomass using a hierarchical modelling approach (Feldpausch *et al*., 2012; Chave *et al*., 2014). Following this approach, it may also be beneficial to model mangroves as a class within global allometric models, rather than developing separate equations with lower sample sizes.

Finally, we note that mangrove soils generally store substantially more carbon than the trees (Hoyos-Santillan, Chavarría, *et al*., 2025). Therefore, whilst we encourage further improvements in allometric equations to constrain aboveground carbon estimates, future research should also focus on improving our estimates of this crucial soil carbon pool.

## Data and code availability

A table summarizing all mangrove trees scanned and used in this analysis is available online. All individual tree point clouds and cylinder models will be openly available on Zenodo

All wood density and carbon content measurements will be openly available on Zenodo

## Acknowledgments

This project was funded by the UK Department for Environment, Food and Rural Affairs (DEFRA) Blue Carbon Fund and the Inter-American Development Bank (IDB) under the project titled “Regional Blue Carbon Monitoring, Reporting and Verification Mechanism” (ATN-BB-19466-RG-T3409).

● TLS data collection in Suriname was funded through the Global Climate Change Alliance (GCCA+ 2) project “ Strengthening the Mangrove Forest Monitoring System”, using a terrestrial laser scanner provided by Ghent University.
● JHS acknowledges the support from the National Audubon Society, the Inter-American Development Bank (“IDB”), and the Smithsonian Tropical Research Institute through the project “Valuing, Protecting and Enhancing Coastal Natural Capital in Panama (PN-T1233) / Blue Natural Heritage”, funded by the UK Blue Carbon Fund of the Department for Environment, Food and Rural Affairs (DEFRA). JHS also acknowledges the support from Panama’s Ministry of the Environment for providing the research permits ARB-012-2022, ARB-002-2022, and ARB-112-2024.
● In Colombia, we especially thank the local mangrove communities and tour operators who made it possible to collect data in the field.

## Author contributions

CRediT: Conceptualization: TDJ, JF, HA; Data curation: TDJ, JF, LLA, JPCG, PCSC, CCMC, AS, JHS, DC, YC, VW, MPO, NSTH, DLA, CR, CP, VMSL, RH, FP, OK, DIRH, MD, HA; Formal Analysis: TDJ; Funding acquisition: FM; Investigation: ; Methodology: TDJ, JF, FJF, CP, KC, HA; Project administration: FM; Resources: ; Software: ; Supervision: ; Validation: ; Visualization: ; Writing – original draft: TDJ, LLA; Writing – review & editing: TDJ, JF, LLA, JPCG, PCSC, CCMC, AS, JHS, EGM, DC, YC, VW, MPO, NSTH, DLA, CR, CP, VMSL, RH, FP, OK, DIRH, TJ, FJF, KC, FM, HA

