## Supplementary materials for "Laser scanning identifies large trees as a major source of uncertainty in mangrove carbon accounting"

#### S1. Measuring the trunk diameter of mangrove trees

The blue text below is 'borrowed' from the Suriname NFI protocol. If this is available online we should simply cite it. We should aim to measure DBH on the point clouds in the same way as they do in the field. The only difference is that we cannot put a tape measure around the tree. We should therefore cut a 2D slice of the point cloud at the relevant point and measure the diameter. Measure in two perpendicular directions and take the average.

The Point of Measurement (POM) for the DBH is taken at a height of 1.3 m. Figure S1 gives a visual description how to measure DBH when the main tree stem shows irregularities (Howard et al., 2014):

- If the tree is fairly straight with a tall trunk the dbh can be measured from the ground parallel to the trunk (Fig S1A)
- If the tree is on a slope, always measure on the uphill side (Fig S1B)
- If the tree is leaning, DBH is taken according to the trees natural height parallel to the trunk (Fig S1C)
- If the tree is forked at or below 1.3 m then measure just below the fork (Fig S1D)
- If the fork is very close to the ground measure as two trees (Fig S1E).
- For trees with tall buttresses exceeding 1.3 m above ground level, stem diameter is usually measured directly above the buttress (Fig S1F). When the height of the buttress becomes a safety issue, estimate the DBH and the new POM.
- For stilt-rooted species (e.g., *Rhizophora* spp.), stem diameter is often measured starting above the highest stilt (Fig S1G). We disregard this measure, when the height of the stilt root becomes a safety issue for the Diameter-taker (person in charge of taking the diameter measurements). In that case, the DBH is taken at the level at which you can visually determine that the bole is uniform. Notice that it is easy to detect where the bole starts tapering and differentiating as a tap root in those trees, go above that until the mentioned criteria is reached, regardless of whether there are more roots above that point. Record the new POM on the field record form.

Keep in mind that when the tree bole becomes difficult to measure DBH (e.g. fused boles) with a diameter tape, use tree calipers to measure the DBH. When using the tree caliper, two measurements are taken per tree stem perpendicular to each other (cross measurement) to compensate for deviations due to the non-circular cross-section and average the two measurements to obtain the DBH.

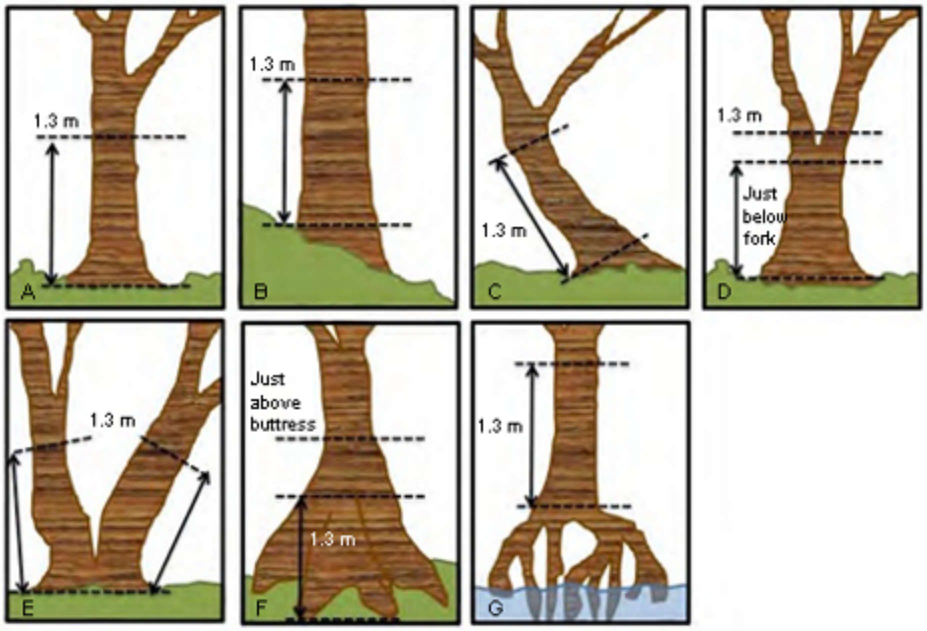


*Figure S1: Estimating diameter at breast height for irregular mangrove trees (modified from Pearson, et al. 2005).*

#### S2. TLS uncertainty in the small branches

Small branches are challenging to accurately model using TLS data. One of the reasons for this is that the laser beam diverges as it travels meaning that returns from the upper canopy have a greater spatial uncertainty. Another reason is simply that these upper canopy branches are less visible from the ground, and TLS does not sample them from above. These sensitivities will vary with the type of scanner and the way in which the data was collected. Our dataset includes Riegl data collected using a regular grid in Suriname and a Leica scanner collected in concentric circles around a focal tree in Colombia, Panama and Jamaica.

In order to test how sensitive our TLS volume estimates were to these small branches, we removed cylinders smaller than 5 cm diameter and re-calculated the total tree volume. This dramatically reduced the number of cylinders in each tree, from a mean of 1786 to 280 per tree. However, removing these cylinders only slightly reduced our estimate of the total woody volume of the trees from a mean of 0.56 m2 to 0.47 m2. This suggests that our biomass estimates are not sensitive to these small branches, and therefore unlikely to be affected by the issues of TLS degradation in the upper canopy. Prop roots were not included in this sensitivity analysis as they were well sampled by the TLS scanning.

The low sensitivity to small branches in this study could be linked to the manual data cleaning that was carried out on the point cloud data. It is impossible to distinguish between branches and leaves in the upper canopy where the TLS data quality is low. We therefore took a conservative approach to the biomass estimates and removed points which we were unsure about. This meant that some small branches were removed from the point cloud during the data cleaning. These noisy areas with low TLS data quality are the regions where large numbers of small branches occur during cylinder fitting. The manual data cleaning therefore reduced this source of uncertainty.

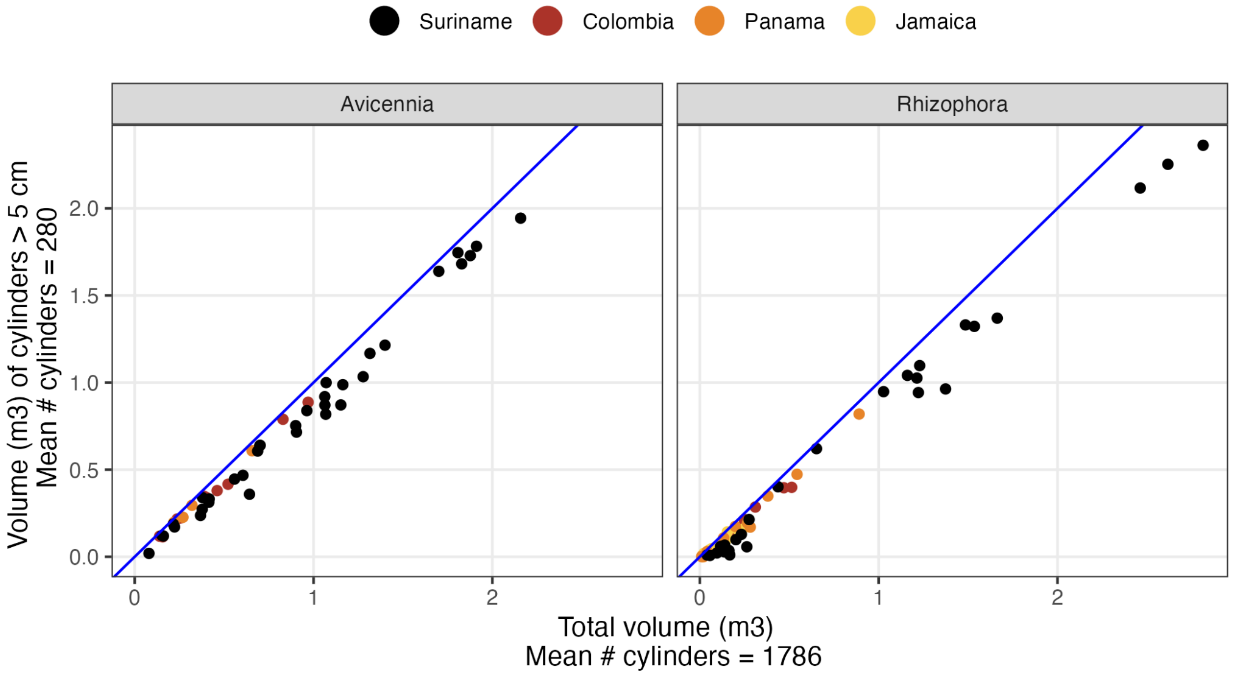


*Figure S2 - Sensitivity of volume estimate to small branches. X-axis is the total volume used in the main text. Y-axis is the volume with small cylinders removed.*

### S3. Comparing cylinder fitting algorithms

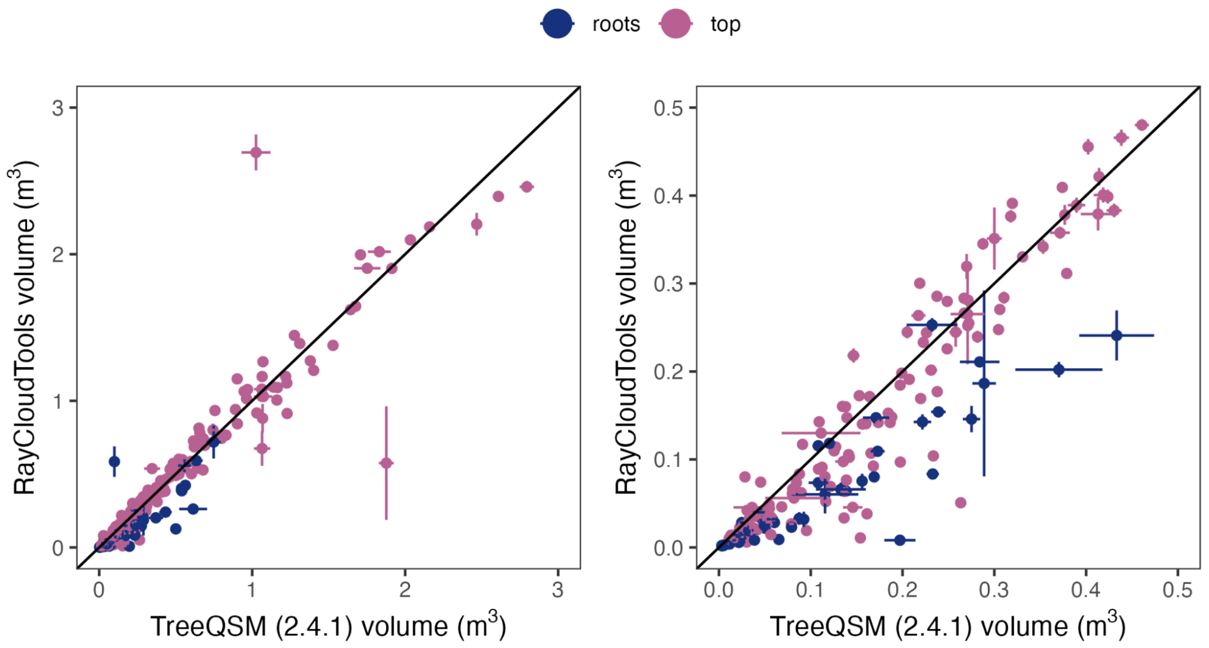


*Figure 3 - Comparison of volume estimates from TreeQSM and RayCloudTools. Left hand panel shows all the data while the right hand panel focuses on the smaller trees.*

It has been suggested that specific cylinder fitting algorithms may overestimate the wood volume. We therefore tested an independent cylinder fitting algorithm (RayCloudTools) in addition to the TreeQSM algorithm used in the main text. We found that volume estimates from the two methods we tested were strongly correlated ($R^2$ = 0.89). We found that TreeQSM volume estimates were on average 0.029 $m^3$ higher than those from RayCloudTools. This strong agreement between the two algorithms gives us confidence that our volume estimates are robust. We note that our data set consists of manually cleaned point clouds. Cylinder fitting algorithms are known to produce unreliable results in point clouds containing large amounts of leaves and noise, so it is likely that the difference between algorithms would be larger in that case.

### S4. Results figures on log-scales

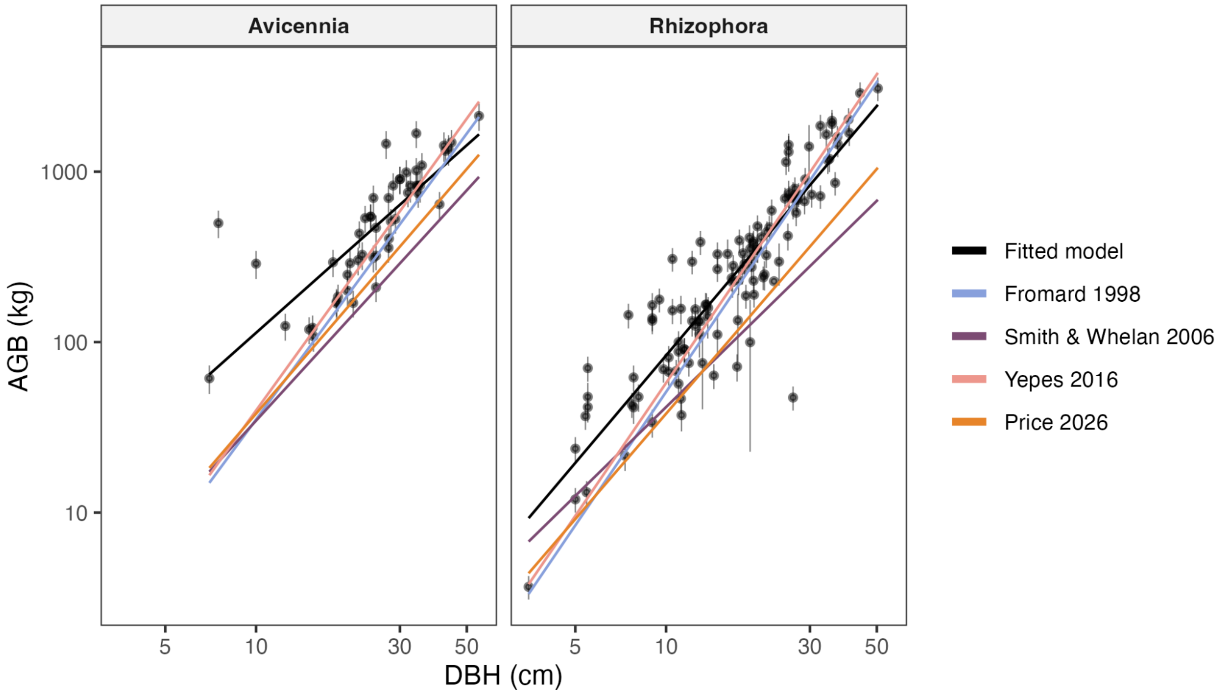


*Figure S4a - Identical to Figure 4 in the main text, except the axes are on log-scales.*

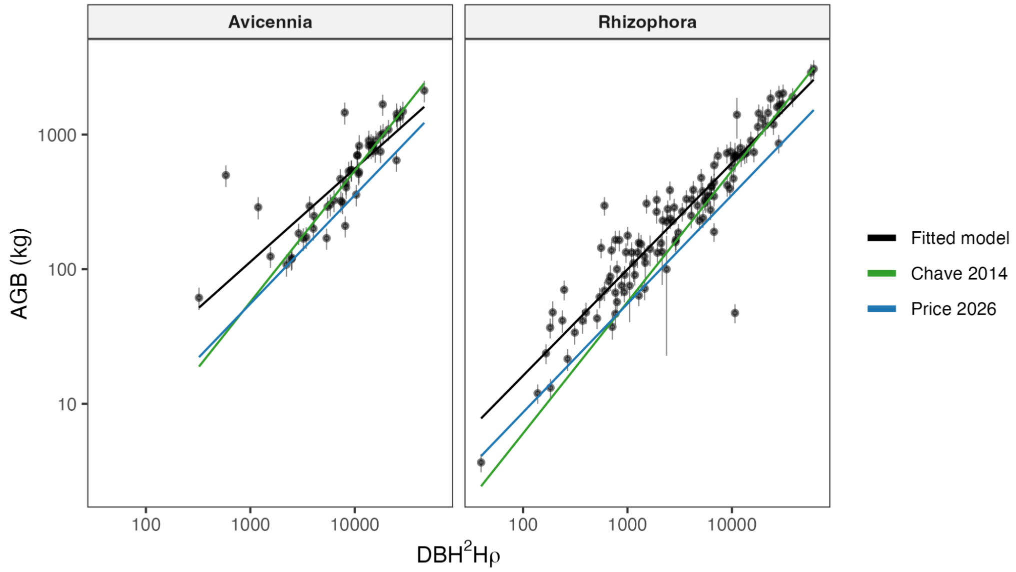


*Figure S5a - Identical to Figure 5 in the main text, except the axes are on log-scales.*

#### S5. Residuals

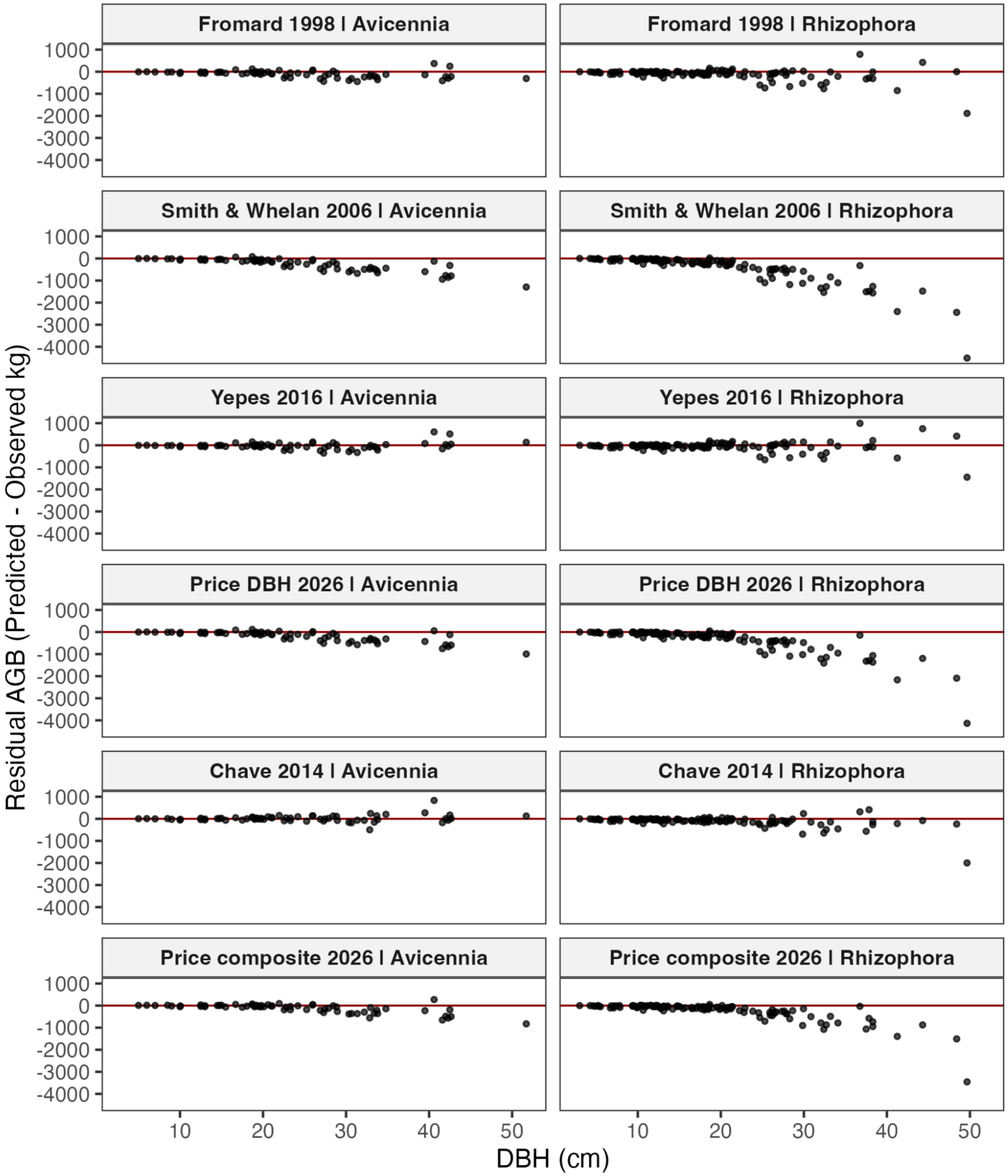


Figure S5 - Residual plots for the different allometric models.

#### S6. Literature review on allometric equations used in mangroves

A systematic literature search was conducted in Web of Science using a Combined Semantic and Boolean Search strategy to identify peer reviewed studies that estimated mangrove biomass in the Americas. The search was performed using the following Boolean expression: (mangrove OR mangroves) AND (biomass OR "aboveground biomass" OR "belowground biomass" OR carbon OR "carbon stocks" OR "carbon estimation") AND ("United States" OR USA OR "United States of America" OR Canada OR Mexico OR "Costa Rica" OR Panama OR Belize OR Cuba OR Jamaica OR "Trinidad and Tobago" OR Colombia OR Brazil OR Brasil OR Ecuador OR Chile OR Peru OR Bolivia OR Paraguay OR Uruguay OR Argentina OR Guyana OR Suriname OR "French Guiana") NOT (soil OR "soil organic carbon" OR sediment OR peat OR "soil carbon" OR "SOC").

Only peer reviewed articles published within the last ten years were included. Reviews, conference proceedings, books, and non research documents were excluded using Web of Science filters. The search was further refined to the following research areas: Forestry, Remote Sensing, Environmental Sciences Ecology, Biodiversity Conservation, Marine Freshwater Biology, Plant Sciences, Geography, Physical Geography, Water Resources, Oceanography, Geochemistry Geophysics, and Science Technology Other Topics.The initial search identified 460 studies.

The first round of manual screening was conducted by reviewing each article in the order returned by Web of Science to identify studies that estimated biomass in mangroves and that used allometric equations. Screening was stopped once a continuous sequence of 30 irrelevant studies appeared when progressing through the ranked results. A second manual screening was then performed; all studies whose abstracts mentioned biomass or aboveground biomass were retained and examined in detail to verify whether they used allometric equations to estimate mangrove biomass. A total of 31 studies met all inclusion criteria. Eligible studies were subsequently classified according to the allometric equations they applied.
